# Glutathione transferase photoaffinity labeling demonstrates GST activation by safeners and NPR1-independent activation by BTH

**DOI:** 10.1101/2023.01.19.524829

**Authors:** Maria Font Farre, Daniel Brown, Maurice König, Brian J. Killinger, Farnusch Kaschani, Markus Kaiser, Aaron T. Wright, Jonathan Burton, Renier A. L. van der Hoorn

## Abstract

Glutathione transferases (GSTs) represent a large and diverse enzyme family involved in detoxification of small molecules by glutathione conjugation in crops, weeds and model plants. Here, we introduce an easy and quick assay for photoaffinity labeling of GSTs to study global GST activation in various plant species. The small molecule probe contains glutathione, a photoreactive group, and a minitag for coupling to reporter tags via click chemistry. Under UV irradiation, this probe quickly and robustly labels GSTs in crude protein extracts. Enrichment and MS analysis of labeled proteins from Arabidopsis identified ten GSTs from the Phi(F) and Tau(U) classes. Photoaffinity labeling of GSTs demonstrated GST activation in wheat seedlings upon treatment with safeners, and in Arabidopsis leaves upon infection with avirulent bacteria. Photoaffinity labeling and proteomics identified six Phi- and Tau-class GSTs that are induced upon treatment with salicylic acid (SA) analog benzothiadiazole (BTH) and these were tested for enhancing immunity in disease assays. Our data confirm that BTH-induced GST activation is independent of NPR1, the master regulator of SA signaling.

## INTRODUCTION

Glutathione transferases (GSTs) are a large and diverse family of enzymes that catalyze the conjugation of the tripeptide glutathione (γGlu-Cys-Gly) to a broad range of small molecule substrates (Sylvestre-Gonon et al., 2019). GSTs have a conserved 3D structure with two conserved pockets: a glutathione binding site (G-site), and a hydrophobic substrate binding site (H-site) (Ding et al., 2017). The G-site is conserved amongst GSTs, stressing the importance of the correct binding and orientation of glutathione for their correct function. The H-site is a more variable pocket, responsible for the promiscuous binding of diverse substrates. Most GSTs are cytosolic and accumulate as soluble homodimers.

In plants, GSTs play important roles in biosynthetic pathways, agrochemical detoxification, plant defence and oxidative stress (Basantani & Srivastava, 2007). Glutathione conjugating GSTs of *Arabidopsis thaliana* include tau (GSTU, 28 members); phi (GSTF, 13 members); theta (GSTT, 3 members); lambda (GSTL, 3 members); and zeta (GSTZ, 2 members) (Dixon et al., 2002; Dixon & Edwards, 2010). GSTU, GSTF, GSTZ and GSTT classes contain a conserved Ser residue in their catalytic site, and can also have peroxidase activity (Sylvestre-Gonon et al., 2019). By contrast, GSTLs are plant-specific Cys-GSTs that contain a conserved catalytic Cys residues and these GSTs reduce small molecules such as β-mercapthoethanol (Dixon & Edwards, 2010). GSTU and GSTF classes are plant-specific and the most abundant and numerous GSTs in plants.

GSTs play a key role in the modification of exogenous compounds (xenobiotics) and their detoxification (Cummins et al., 2011). In agrochemical detoxification, GSTs are responsible for conjugating glutathione to herbicides, converting them into hydrophilic compounds with reduced mobility and toxicity. Metabolic herbicide resistance in weeds is associated with elevated levels of herbicide-catabolizing enzymes, including GSTs. In blackgrass (*Alopecurus myosuroides*) and ryegrass (*Lilium* spp.), the upregulation of GSTs is associated with resistance to multiple herbicides in resistant field populations, with *Am*GSTF1 commonly upregulated in resistant blackgrass populations (Cummings et al., 2013; Dücker et al., 2019, 2020; Tetard-Jones et al., 2018). Other examples are the high constitutive expression of GSTF2, associated with atrazine resistance in two waterhemp (*Amaranthus tuberculatus*) populations, and high GST levels in resistant populations of American sloughgrass (*Beckmannia syzigachne*) (Evans et al., 2017; Pan et al., 2016; Wang et al., 2020). Furthermore, safeners improve herbicide tolerance in crops by selectively inducing herbicide metabolizing activities in cereal crops but not in weeds (Hatzios & Burgos, 2003; Jablonkai, 2013). For instance, safener isoxadifen-ethyl reduces maize injury caused by herbicide nicosulfuron, associated with the induction of GSTs (Sun et al., 2017). Similarly, safeners fenclorim and metcamifen protect rice against injury by pretilachlor an clodinafop herbicides, respectively, again associated with GST induction (Brazier-Hicks et al., 2020; Hu et al., 2020; 2021; Taylor et al., 2013).

GSTs are also induced during biotic stress and play a role in plant defense (Gullner et al., 2018; Nianiou-Obeidat et al., 2017). Salicylic acid treatment induces the expression of phi and tau class GSTs in Arabidopsis and tomato (Sappl et al., 2004; Csiszar et al., 2014). A study on maize revealed a genetic linkage between a GST-encoding gene and multiple disease resistance to three different fungal leaf pathogens (Wisser et al., 2011). Likewise, silencing of *Nb*GSTF1 in *Nicotiana benthamiana* increased susceptibility to *Colletotrichum* infections (Dean et al., 2005).

The different roles of GSTs in plants are not yet fully understood and this research field is hampered by the large number and diversity of GSTs. Here, we establish photoaffinity labeling of GSTs as an easy way to detect GSTs in various tissues of model plants, crops and weeds. We took advantage of a glutathione-based photoaffinity probe that successfully labeled recombinant human GSTs and endogenous GSTs from mouse liver and lung tissues (Stoddard et al., 2017). This photoaffinity probe contains glutathione; a benzophenone photoreactive group; and an alkyne minitag that can be labeled with a fluorophore of biotin via click chemistry. We optimized labeling parameters, identified the probe targets and studied GSTs in plants responding to agrochemicals and biotic stress.

## RESULTS

To explore GST profiling in plants, we studied the chemical reactivity of the glutathione-based photoaffinity probe DB478 (**Figure 1A**), which is a re-synthesized replica of GSTABP-G (Stoddard et al., 2017). DB478 consists of three parts: i) a glutathione moiety, which directs DB478 toward the glutathione binding site (G-site) in GSTs; ii) a benzophenone photoreactive group that facilitates the covalent crosslinking of the probe to a C-H bond in a nearby amino acid upon UV irradiation; and iii) an alkyne minitag for subsequent coupling to a fluorescent or affinity reporter tag via ‘click chemistry’ (**Figure 1B and 1C**).

**Figure 1.**
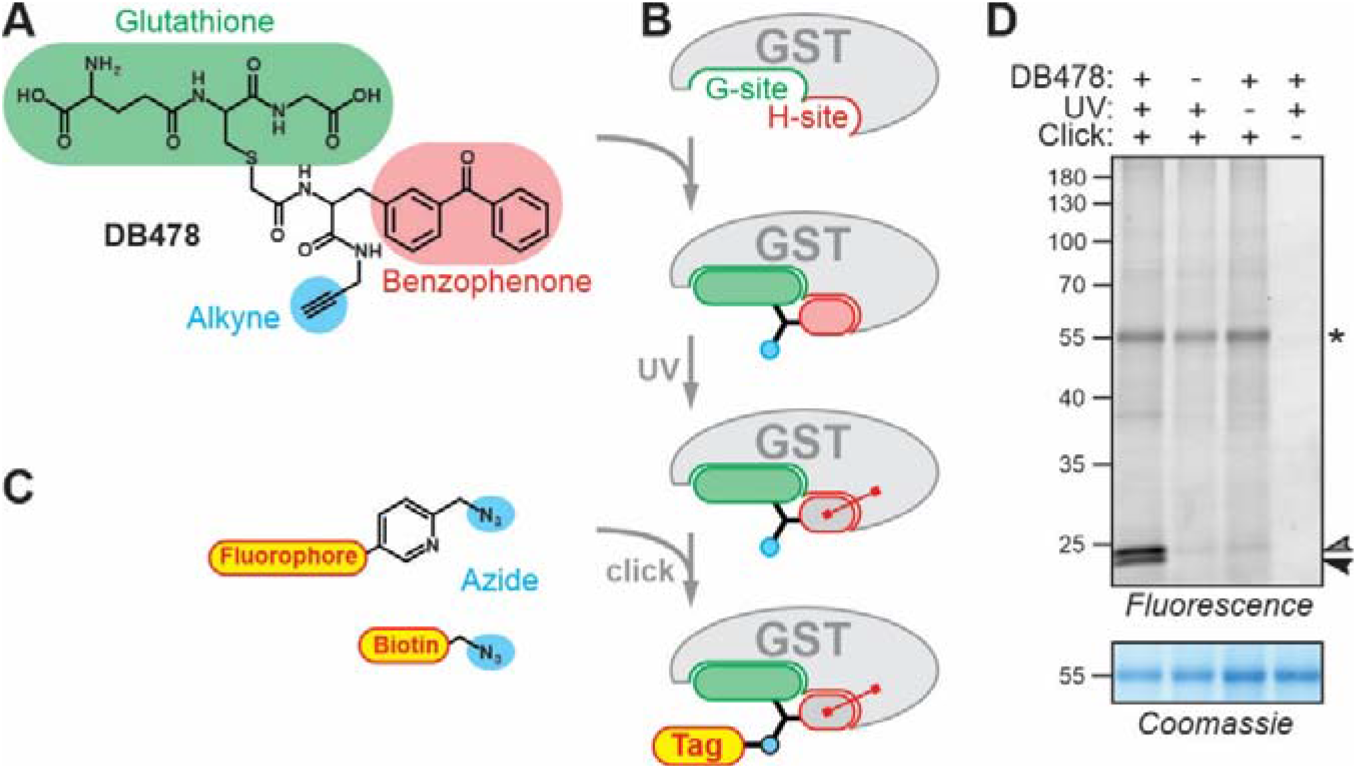
Photoaffinity labeling of GSTs. **(A)** Structure of the GST photoaffinity probe DB478. **(B)** Procedure of photoaffinity labeling of GSTs. **(C)** Structures of the used reporters. **(D)** Photoaffinity labeling of Arabidopsis leaf extracts requires probe, UV light and click chemistry. An Arabidopsis leaf extracts was incubated with and without 5 μM probe and irradiated with UV light for 45 minutes, and coupled to a fluorophore via click chemistry. Fluorescent proteins were visualized by in-gel fluorescence scanning. Coomassie staining is show as loading control. Arrowheads: Specific labeling of GSTs; *: aspecific labeling of the large subunit of rubisco (RbcL).

### Photoaffinity labeling of leaf extracts displays 23-24 kDa signals that depend on conditions

To study GST photoaffinity labeling, Arabidopsis leaf extracts were incubated with and without 5 μM DB478, irradiated with UV at 365nm, and coupled to a fluorophore with click chemistry. Fluorescence scanning of the labeled extracts separated on protein gels revealed two signals at 23-24 kDa, consistent with the molecular weight (MW) of GSTs (**Figure 1D**). These 23-24 kDa signals are absent in the no-probe-control, and requires both UV treatment and click chemistry (**Figure 1D**). Another, weaker signal at 55 kDa is caused by non-selective coupling of the reporter tag probably to rubisco as it does not require DB478 or UV exposure, but depends on click chemistry (**Figure 1D**).

We next characterized GST photoaffinity labeling further by studying the effect of probe concentration, UV exposure time, pH and presence of GST inhibitors. These experiments demonstrate that DB478 labeling reaches saturation at 5μM (**Figure 2A**), and that a 30-minute UV exposure is sufficient to detect robust DB478 labeling (**Figure 2B**). DB478 labeling is strongly dependent on pH, with poor labeling at acidic pH, and increased labeling at neutral to basic pH (**Figure 2C**). The two detected signals at 23-24 kDa are not differentially labeled at various probe concentrations, UV exposure times or pH. Therefore, the remaining experiments were performed with 5 μM DB478, 45-minute UV irradiation and at pH 7.5, which mimics the cytonuclear pH where GSTs reside.

**Figure 2.**
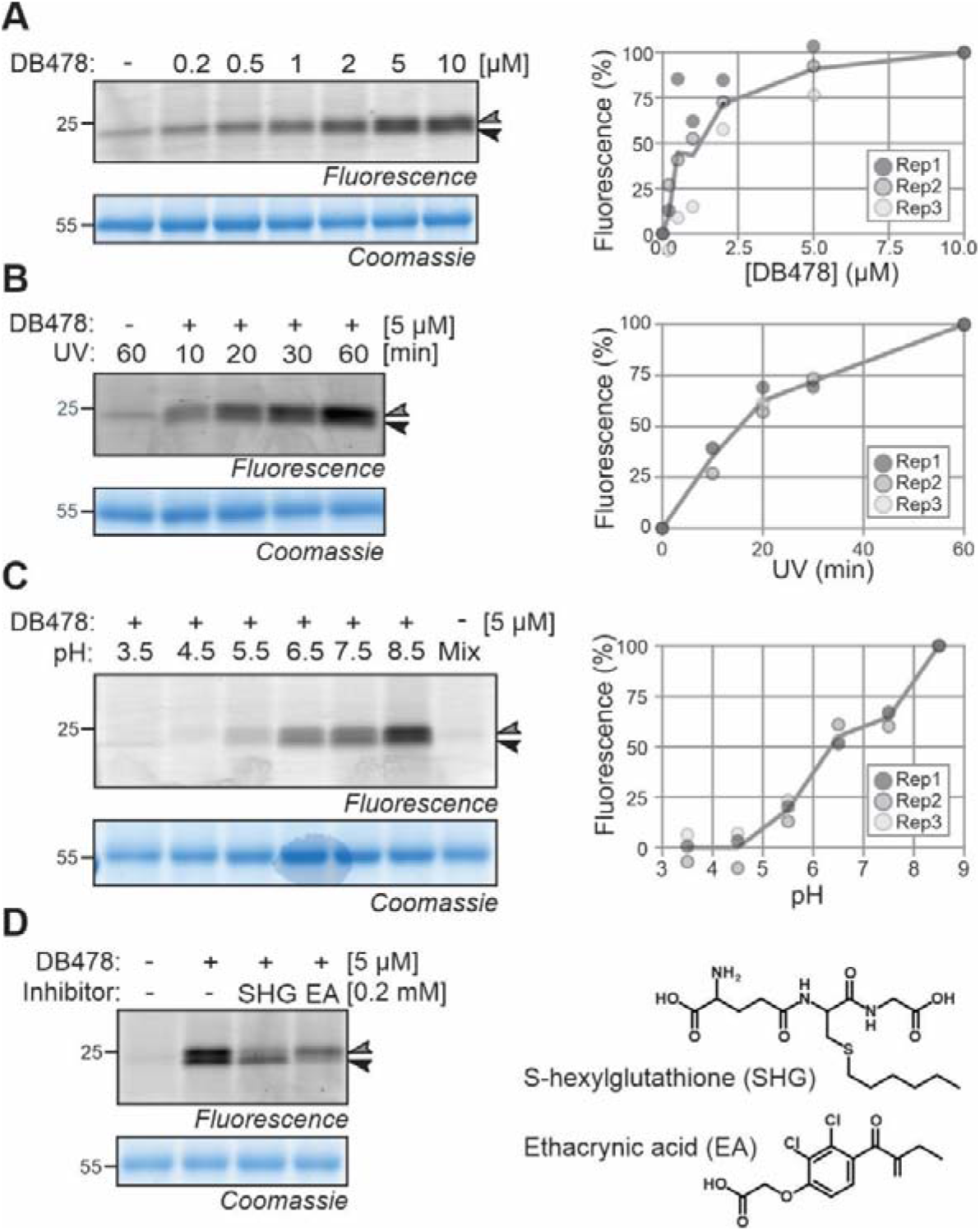
GST labeling depends on conditions. **(A)** Photoaffinity labeling saturates at 5 μM probe concentration. Arabidopsis leaf extracts were incubated with various probe concentrations and exposed to UV for 30 minutes. **(B)** Photoaffinity labeling depends on requires >30 minutes UV treatment. Arabidopsis leaf extracts were incubated with 5 μM probe and exposed to UV for various times. **(C)** Photoaffinity labeling requires pH>6. **(D)** GST inhibitors differentially suppers photoaffinity labeling. Arabidopsis leaf extracts were incubated with 5 μM probe at various pH and exposed to UV for 30 minutes. Samples were further analyzed as described in **Figure 1D**. Plotted are signal intensities from three independent replicates normalized to the signal with highest intensity.

We next confirmed GST labeling by competing DB478 labeling with known GST inhibitors. We tested S-hexylglutathione (SHG), a glutathione analogue competing for the G-site of GSTs, and ethacrynic acid (EA), a substrate/inhibitor for the H-site of GSTs (Allocati et al., 2018). Interestingly, pre-incubation of Arabidopsis leaf extracts with SHG suppressed labeling of the top signal, whereas preincubation with EA suppressed labeling of the bottom signal (**Figure 2D**), indicating that these signals are caused by different GSTs that have contrasting sensitivities to SHG and EA inhibitors.

### GSTs are the main targets of DB478

To identify the targets of DB478, Arabidopsis leaf extracts were incubated with 5 μM DB478 and labeled proteins were coupled to a biotin affinity tag for subsequent enrichment on streptavidin beads. Purified proteins were on-bead digested with Lys-C and trypsin and digested peptides were identified by liquid chromatography tandem mass spectrometry (LC-MS/MS). The pull down was performed three times for DB478 labelled samples and three times for click control samples that followed the same procedure except no DB478 was added.

After removing contaminants and retaining only proteins that were detected in all three samples of either the probe or the no-probe control, a total of 2205 proteins were identified. Of the 29 proteins that were significantly enriched in the probe-labeled samples, we identified nine GSTs from two different classes (Phi(F) and Tau(U), **Figure 3A**). The highest enrichments (log2FC>6) and highest MS signal intensities are all caused by three members of the Phi class (GSTF8, GSTF9, and GSTF10) and two members of the Tau class (GSTU7 and GSTU16). We also detected GSTF2/3, GSTF6, GSTF7 and GSTU13 with high MS signal intensities but with lower but significant enrichments (log2FC>2). All ten significantly enriched GSTs were detected with many unique peptides and 50-85% sequence coverage and have a predicted MW of 23.5-29.2 kDa (**Figure 3B**), consistent with the fluorescent signals seen from protein gels. The other enriched proteins were detected at much lower MS intensities and do not have a predicted MW of 23-30 kDa. The majority these other proteins were unchanged between the DB478 and click control (grey dots in **Figure 3A**) and these include endogenously biotinylated proteins BCCP, MCCA and ACC1 as well as abundant proteins such as the large and small subunits of rubisco (RbcL and RbcS, respectively). We consider them being caused by click chemistry background labeling, consistent with the background labeling seen on fluorescent gels (**Figure 1D**).

**Figure 3.**
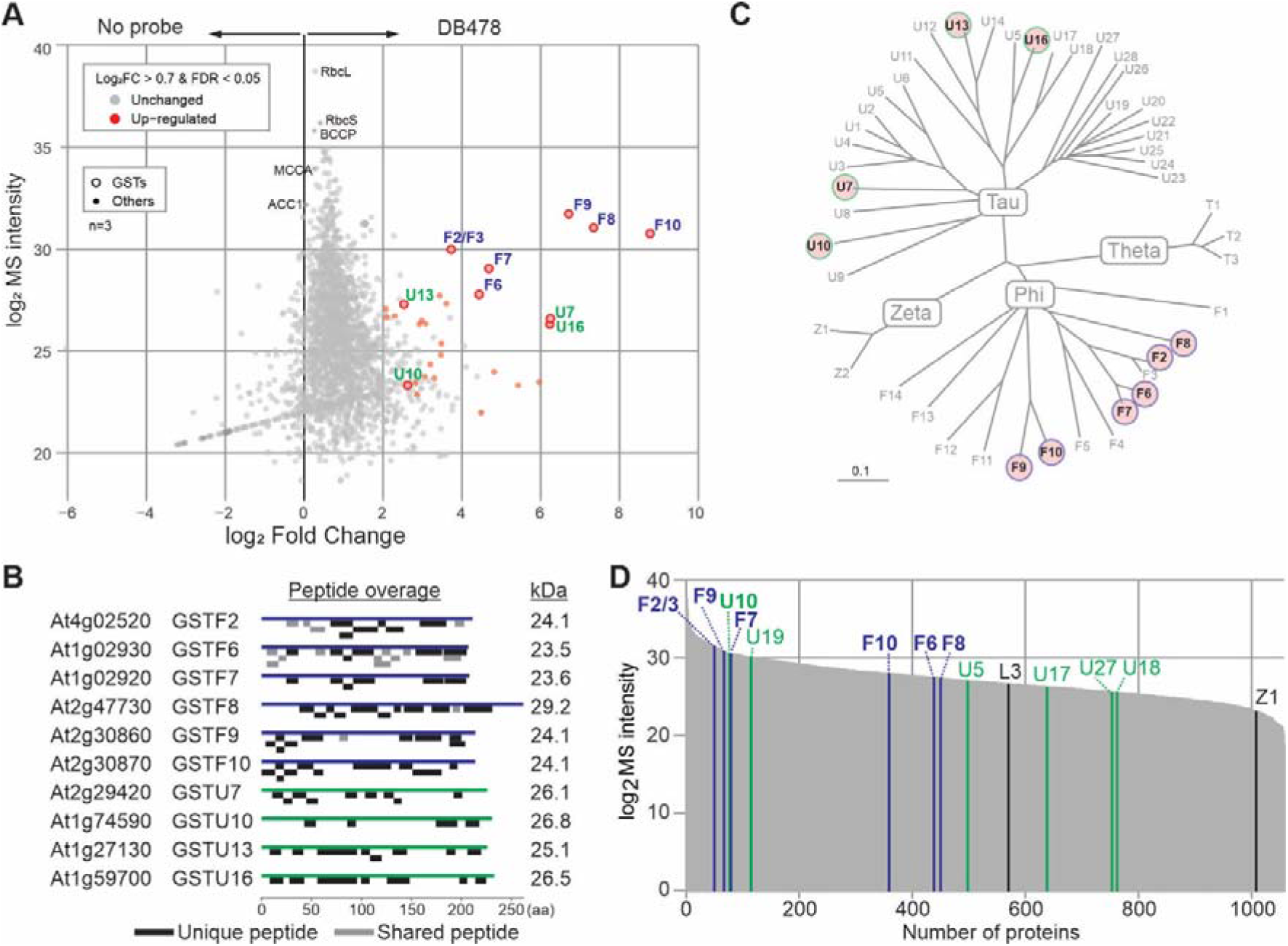
Proteomics confirms GST enrichment upon labeling. **(A)** GSTs are enriched in probe-labeled samples. Arabidopsis leaf extracts were labeled with and without 5 μM probe, crosslinked and coupled to biotin via click chemistry and enriched on streptavidin beads. Beads were treated with trypsin and released peptides were analyzed by MS. Plotted are the MS protein intensities plotted against the fold change for all proteins that were detected in all three replicates of the probe labeled samples. Highlighted are enriched (red), all detected GST proteins (circles), abundant non-enriched proteins (RbcL, RbcS) and endogenously biotinylated proteins BCCP, MCCP and ACC1. **(B)** High protein coverage of GSTs. Unique and shared peptides identified by proteomics (A) were mapped onto the GST protein sequences. The predicted molecular weight in kDa for each GST is on the left. **(C)** Identified GSTs highlighted in phylogeny of Arabidopsis GSTs. Tree is adapted from (Wagner et al., 2002). Unrooted bootstrapped tree (n = 5000) is based on a multiple sequence alignment by ClustalX of the full-length protein sequences of all Arabidopsis GSTs. **(D)** In-solution digest (ISD) of the same Arabidopsis leaf extracts used for labeling. Shown are the proteins detected in all n=3 replicates, ranked on their average LFQ intensity, with the detected GSTs highlighted.

Although we only identified GSTs from the Tau and Phi class, the identified GSTs distribute well over the phylogenetic tree of Arabidopsis GSTs (**Figure 3C** Wagner et al., 2002), indicating that DB478 can label diverse GSTs. In-solution-digests of the proteomes used for labeling showed that all five Phi-class GSTs that were detected in the proteome were also detected upon labeling and enrichment (**Figure 3D**). By contrast, except for GSTU10, five other members of the Tau class are present in the proteome but not enriched upon labeling (GSTU5, -U17, -U18, -U19, -U27), indicating that not all GSTUs can be labeled by DB478. Conversely, we detected three Tau-class GSTs (GST-U7, -U13 and -U16) upon labeling that were not detected in the proteome, indicating their efficient enrichment upon labeling.

### GST photoaffinity labeling of other plant species

To profile GSTs in other plant species, we tested leaf extracts of *Nicotiana benthamiana*, an important model plant used frequently for transient expression experiments (Bally et al., 2018). However, only weak signals were detected, in stark contrast to Arabidopsis leaf extracts (**Figure 4A**). Leaf extracts of *N. benthamiana* usually turn brown quickly during labeling (**Figure 4B**) in contrast to Arabidopsis leaf extracts. Browning is presumably caused by the oxidation of polyphenols by polyphenol oxidase (PPO) activity, which is frequently observed in plant extracts (Hamdan et al., 2022). Oxidation also results in crosslinking of proteins, which severely hampers their separation on protein gels, consistent with a strongly reduced RbcL signal from the Coomassie-stained gel (**Figure 4A**). Protein oxidation can be prevented by performing the extraction in 5 mM β-mercaptoethanol (βME), resulting in well-resolved RbcL signals on Coomassie-stained gels (**Figure 4A**). Importantly, in the presence βME, labeling of *N. benthamiana* leaf extracts with DB478 results in a specific 25 kDa signal, consistent with GST labeling (**Figure 4A**).

**Figure 4.**
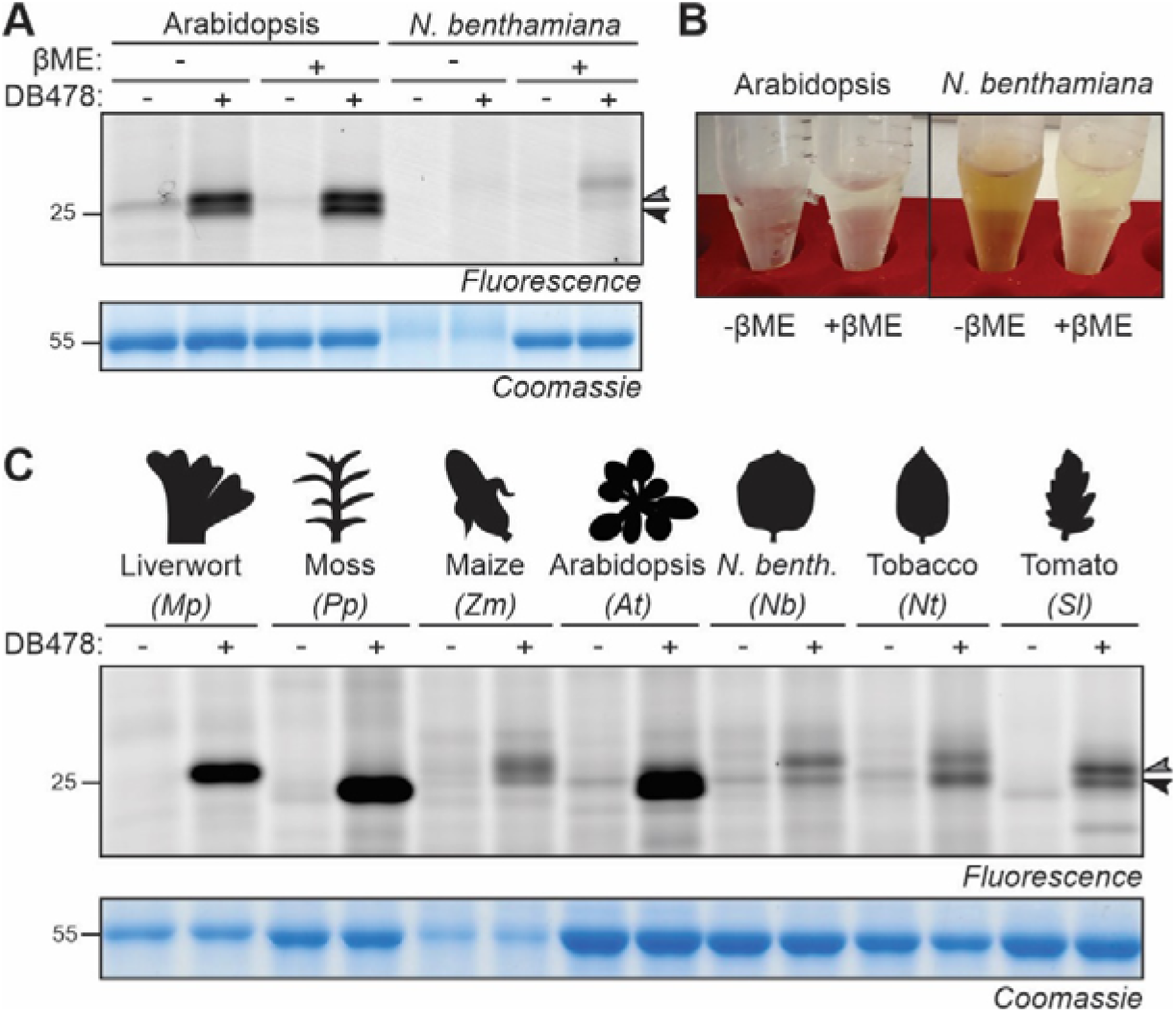
GST labeling of other plant species. **(A)** Photoaffinity labeling of GSTs in *N. benthamiana* requires extraction with reducing agent. Leaf extracts of Arabidopsis and *N. benthamiana* were generated with and without 3 mM β-mercaptoethanol (βME); incubated with and without 5 μM probe and exposed to UV light; and visualized by fluorescence scanning and coomassie staining. **(B)** βME avoids sample browning during labeling. **(C)** GST labeling in leaf extracts of various plant species in the presence of βME. Leaf extracts were generated with 3 mM βME; incubated with and without 5 μM probe and exposed to UV light; and visualized by fluorescence scanning and coomassie staining.

We next tested if DB478 could label GSTs in additional plant species. We produced leaf extracts of two nonvascular plants (the livewort *Marchantia polymorpha* and the moss *Physomitrella patens*) and five angiosperms (the monocot *Zea mays* and four eudicots: *Arabidopsis thaliana*, *Nicotiana benthamiana*, *Nicotiana tabacum* and *Solanum lycopersicum*). Labeling of leaf extracts with DB478 in the presence of βME consistently resulted in signals in the 25 kDa region, consistent with GST labeling (**Figure 4C**). These signals vary in intensity and MW, reflecting with the diversity of GSTs in various plant species.

### Safener treatment of wheat increases GST labeling

GSTs are important detoxifying enzymes that contribute to herbicide resistance (Jablonkai, 2013). Having established GST profiling on various plants, we investigated the effect of herbicides and safeners on GSTs. Fenoxaprop-P-ethyl is an extensively used post-emergence herbicide against grass weeds in wheat crops, usually applied in combination with safeners such as mefenpyr-diethyl (**Figure 5A**, Taylor et al., 2013). To test whether the safener treatment activates GSTs, we incubated four-day-old wheat seedlings in liquid medium containing 100 μM safener (S), or herbicide (H), or both (S+H), or neither (C) for one day (**Figure 5B**). Subsequent DB478 labeling revealed that seedlings treated with safener alone (S) of with safener and herbicide (S+H), has significantly elevated GST labeling, whereas seedlings treated with herbicide (H) have GST labeling that is similar to the control (C) (**Figure 5C**). A possible synergistic effect of the dual S+H treatment was seen in various experiments but is not statistically significant (**Figure 5C**). This experiment illustrates that DB478 labeling can be used to study the activation of GSTs upon agrochemical treatment, probably in both weeds and crops.

**Figure 5.**
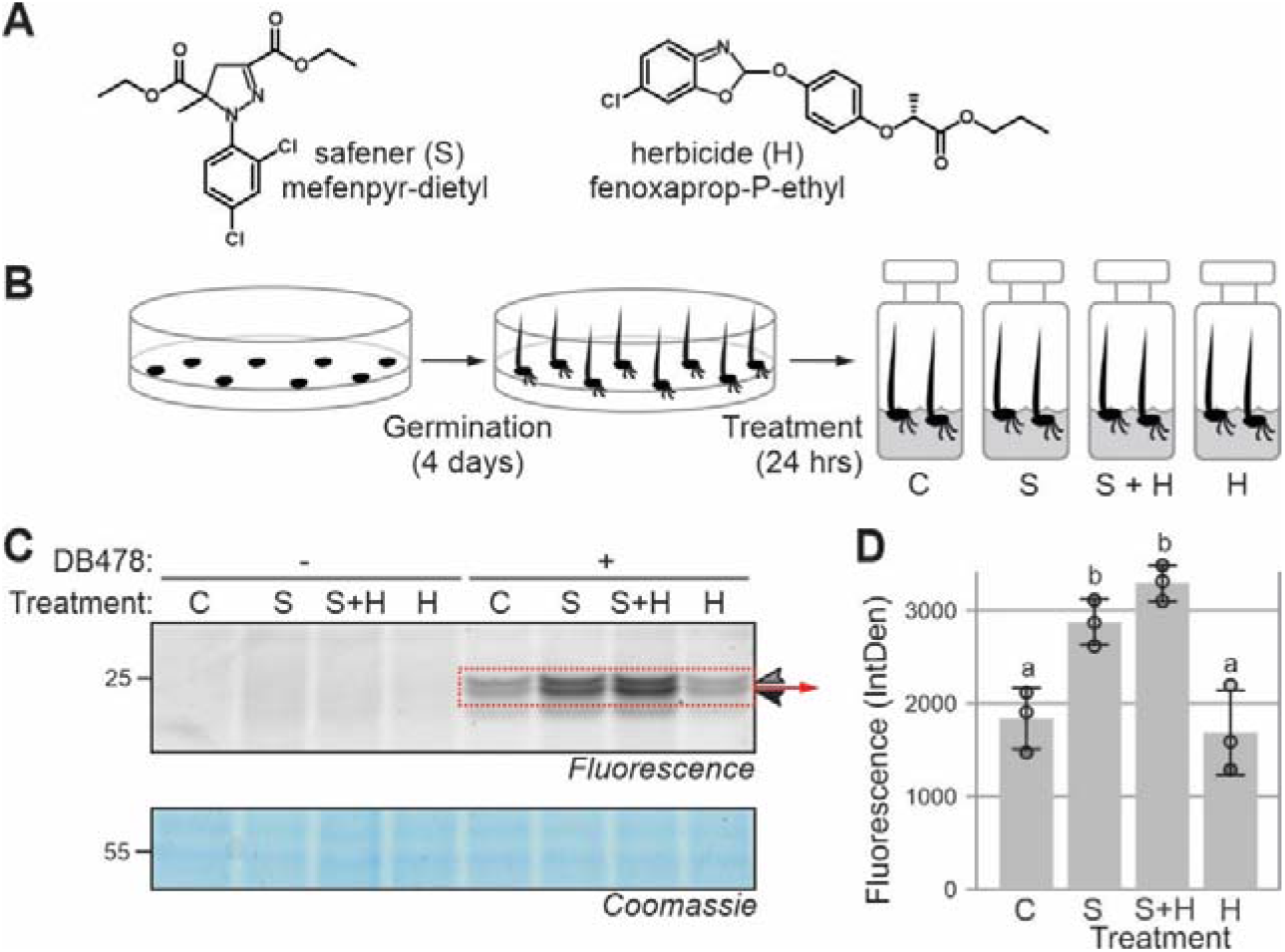
Safener activates GSTs in wheat. **(A)** Structures of safener and herbicide used in this experiment. **(B)** Experimental procedure of agrochemical treatment. **(C)** Safener treatment activates GST activities. Leaf extracts of seedlings incubated with/out safener and/or herbicide were labeled with DB478 and fluorescently labeled proteins were detected from protein gels by fluorescence scanning and quantified. Coomassie staining is shown as a loading control. **(D)** Quantified fluorescence of the 22-24 kDa signals from n=3 biological replicates, analysed by ANOVA. Groups a and b are statistically different from each other (p<0.05). Error bars represent standard deviation of n=3 biological replicates.

### Biotic stress and BTH activate GSTs

To study GST activation during biotic interactions, we infiltrated Arabidopsis leaves with wild-type *Pseudomonas syringae* pv. *tomato* DC3000 (*Pto*DC3000), which is pathogenic on Arabidopsis, and *Pto*DC3000 expressing avrRpt2 and avrRpm1, which are both avirulent on the Arabidopsis *Col-0* ecotype because this carries both *RPS2* and *RPM1* resistance genes, respectively. Interestingly, whilst wild-type *Pto*DC3000 had minimal impact on GST photoaffinity labeling, GST labeling is clearly elevated upon infection with both avirulent strains (**Figure 6A**).

**Figure 6.**
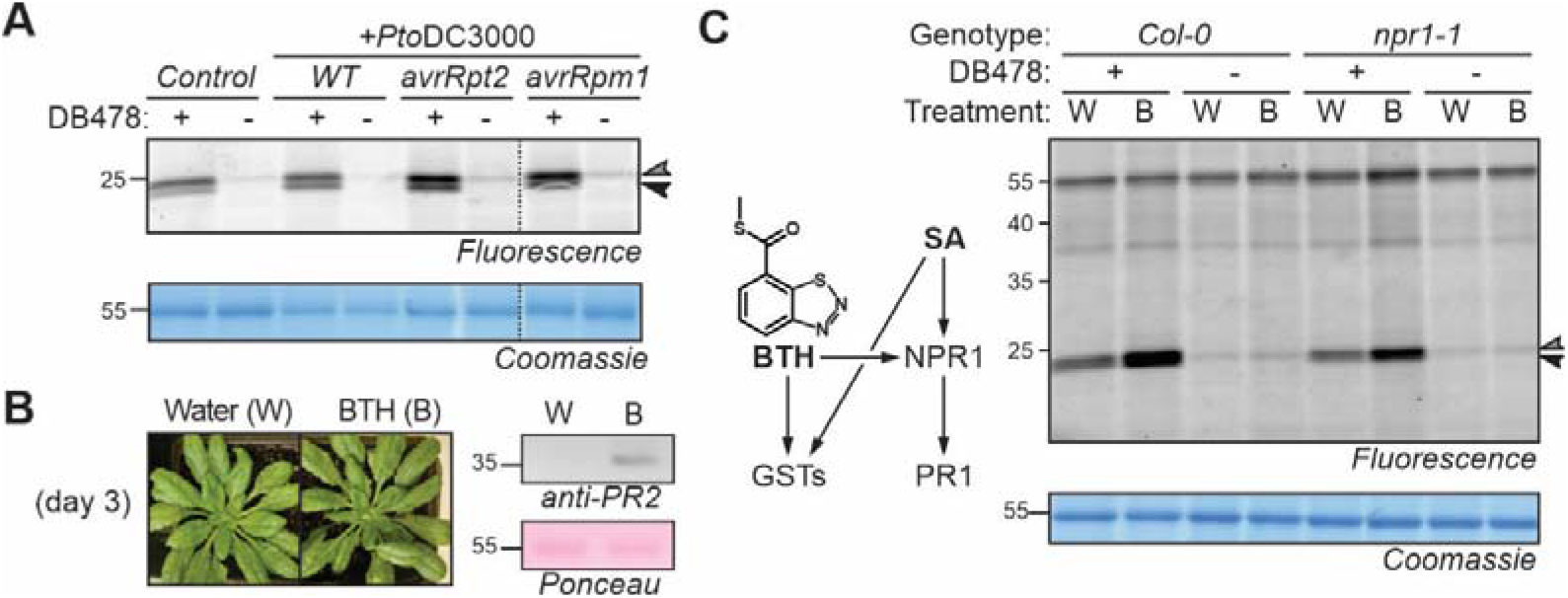
Pathogens and BTH activate GSTs independently of NPR1. **(A)** Avirulent *P. syringae* causes increased GST labeling in Arabidopsis. Arabidopsis leaves were infiltrated with *Pto*DC3000 wild-type (WT) or carrying plasmids encoding avrRpt2 or avrRpm1. Total extracts generated at day-2 were labeled with DB478, coupled to a fluorophore and analysed by fluorescent scanning of protein gels. **(B)** BTH treated plants. Adult Arabidopsis plants were watered with and without BTH for 3 days and proteins were extracted and analyzed by western blot using anti-PR2 antibodies and Ponceau staining. **(C)** BTH-induced GST labeling is not dependent on NPR1. The samples of *Col-0* wild-type and *npr1-1* mutant plants treated with water/BTH for 3 days were labeled with 5 μM DB478, coupled to a fluorophore with click chemistry, followed by in-gel fluorescence scanning and Coomassie staining.

Since many responses triggered by avirulent bacteria are dependent on stress hormone salicylic acid (SA), we tested if treatment with benzothiadiazole (BTH), which is a mimic of SA, would be sufficient to induce GST labeling. BTH treatment for 3 days triggers PR2 protein accumulation without drastically reducing growth (**Figures 6B**). GST photoaffinity labeling of these samples showed a robust and significant increase of GST labeling in BTH treated samples when compared to the water-treated control samples (**Figure 6C**).

Gene induction induced by BTH and avirulent pathogens involves the salicylic acid (SA) signaling pathway. Activation of many SA-responsive genes requires NPR1 (nonexpressor of PR1), which acts downstream of SA by activating TGA transcription factors that bind to *activator sequence-1* (*as-1*) promoter elements of SA-responsive genes (Kumar et al., 2022). The *as-1* elements are also present in promotors of GST-encoding genes (Blanco et al., 2005). We therefore tested if BTH-induced GST labeling requires NPR1. Notably, increased GST labeling upon BTH treatment also occurs in the *npr1-1* mutant (**Figure 6C**), demonstrating that NPR1 is not required for the increased GST labeling in samples from BTH-treated plants.

### BTH-induced GSTs do not increase immunity in transient assays

To investigate BTH-induced GSTs further, we labelled total leaf extracts for Arabidopsis treated with water and BTH with DB478 and purified and identified labelled proteins by mass spectrometry. Amongst the significantly BTH-induced proteins we detected GSTU9 and U10 and GSTF2/3, F6 and F7, whilst GSTU4 just dropped below significance level (**Figure 7A**). Many additional proteins were detected at lower intensities, presumably caused by nonspecific binding. This includes Pathogenesis Related-1 (PR1), which was detected as significantly induced upon BTH treatment (**Figure 7A**), consistent with being a marker protein for SA signaling (Zhang et al., 1999).

**Figure 7.**
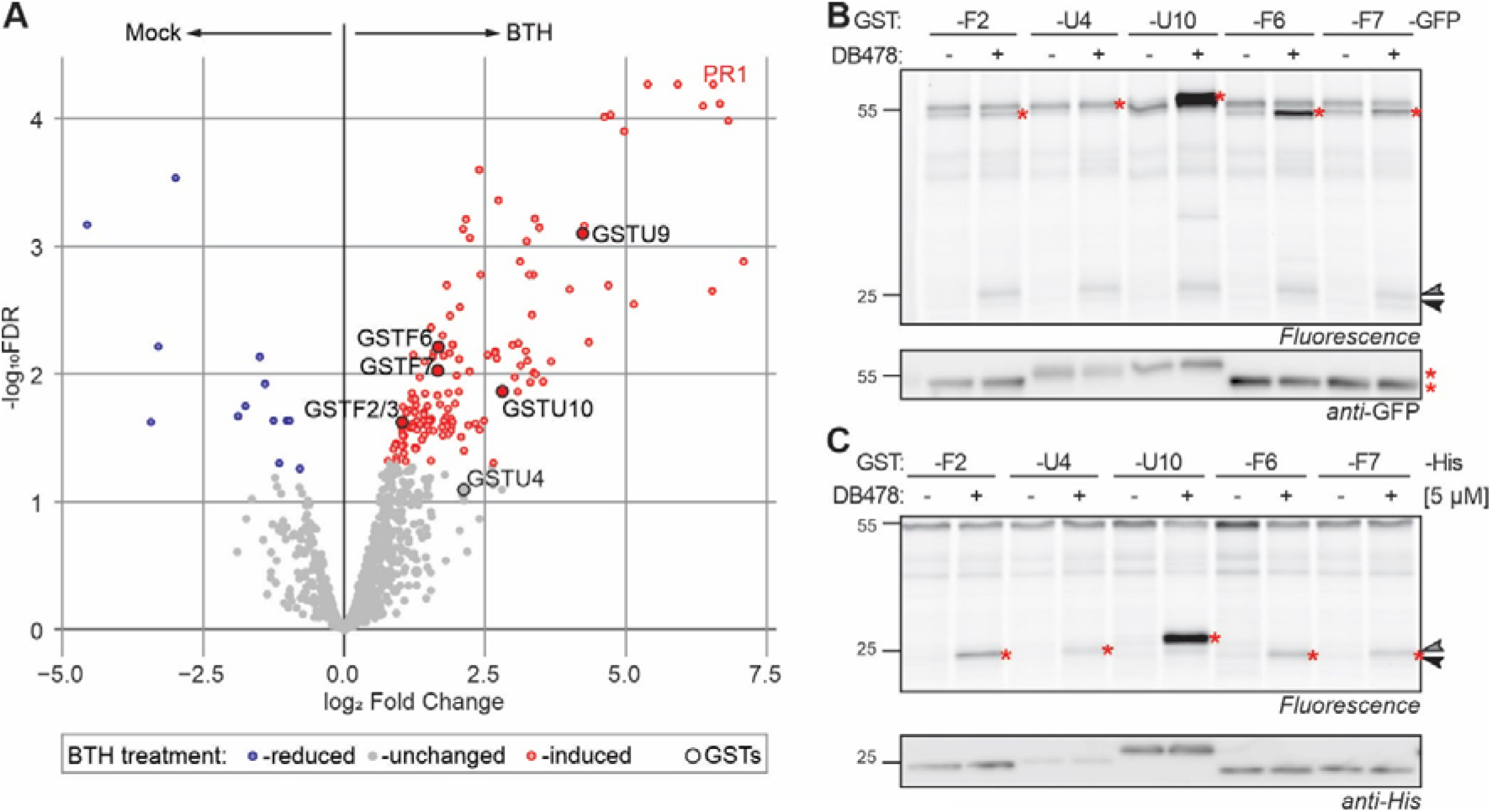
Transient overexpression of BTH-induced GSTs **(A)** BTH-induced GSTs. Leaf extracts from water- and BTH-treated Arabidopsis plants were labelled with DB478 in triplicate. Labeled proteins were coupled to biotin-azide via click chemistry and enriched on streptavidin beads. On-bead digested proteins were analyzed by MS and plotted in volcano plots where log2 of fold change (log2FC) was plotted against their significance (-log_10_FDR) for n=3 biological replicates. Proteins were significantly reduced (blue) induced (red) or unchanged (grey) upon BTH treatment. GSTs and PR1 are highlighted. **(B)** Labeling of GST-GFP fusion proteins upon agroinfiltration. Arabidopsis GSTs were fused to a C-terminal GFP tag and expressed in *N. benthamiana* by agroinfiltration. Extracts of agroinfiltrated leaves taken at 4 days upon agroinfiltration were labeled with and without5 μM DB478, coupled to a fluorophore via click chemistry, and visualized by in-gel fluorescence scanning. Fusions were detected by western blot with anti-GFP antibody. Highlighted are labeled GFP-tagged GSTs (*) and endogenous GSTs (arrowheads). **(C)** Labeling of GST-His fusion proteins upon agroinfiltration. Arabidopsis GSTs were fused to a C-terminal His tag and expressed in *N. benthamiana* by agroinfiltration. Extracts of agroinfiltrated leaves taken at 4 days upon agroinfiltration were labeled with and without5 μM DB478, coupled to a fluorophore via click chemistry, and visualized by in-gel fluorescence scanning. Fusions were detected by western blot with anti-GFP antibody. Highlighted are labeled GFP-tagged GSTs (*) and endogenous GSTs (arrowheads).

To investigate the six BTH-induced GSTs further, we cloned them into agroinfiltration vectors with C-terminal His and GFP tags. While cloning of GSTU9 failed, the other five GSTs were successfully expressed by agroinfiltration of *Nicotiana benthamiana*, resulting in the detection of His-tagged and GFP-tagged fusion proteins (**Figure 7B** and **7C**). DB478 labelling of these samples showed clear labelling of all these GSTs, and strong labeling of GSTU10 and GSTF6 much beyond the level of endogenous GSTs, detected at 23-24 kDa (**Figure 7B** and **7C**).

To test if these defence-induced GSTs contribute to immunity, we challenged leaves transiently over expressing GFP-tagged GSTs with pathogens (**Figure 8A**). Spray infection with *Pto*DC3000 and monitoring bacterial growth in the ‘agromonas’ assay (Buscaill et al., 2021), revealed no altered bacterial growth in 3 days, in contrast to the tomato FLS3 positive control (**Figure 8B**). Likewise, also droplet-inoculation with zoospores of the oomycete pathogen *Phytophthora infestans* did not alter lesion size in tissues expressing GST-GFP proteins, despite the clear GFP fluorescence detected in agroinfiltrated tissues (**Figure 8C**). We conclude that GSTF2, -F6 and -U10 to not increase immunity to *Pto*DC3000 and *P. infestans* in these transient disease assays.

**Figure 8.**
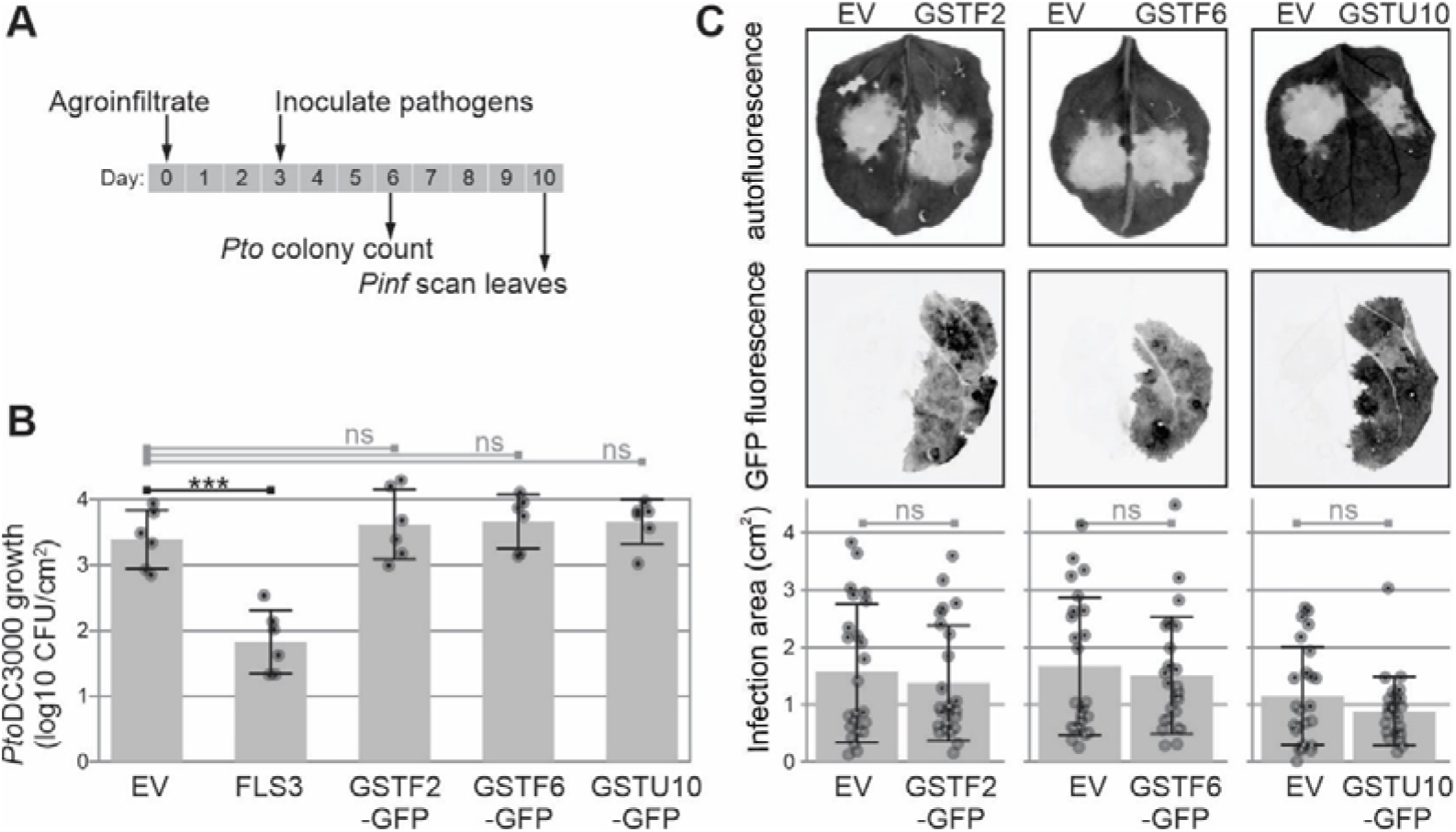
Three BTH-induced GSTs do not increase immunity in transient assays. **(A)** Experimental procedure for detecting roles of GSTs in immunity. Three days after agroinfiltration, agroinfiltrated leaves were spray-infected with *Pto*DC3000 or droplet-inoculated with *P. infestans*. Pathogen growth is determined 3 and 5 days later for *Pto*DC3000 and *P. infestans*, respectively. **(B)** Transient expression of GSTs does not impact *Pto*DC3000 growth. Agroinfiltrated leaves expressing empty vector (EV), FLS3, GST-F2-GFP, GSTF6-GST or GSTU10-GFP were spray-inoculated with *Pto*DC3000 at OD=1. Bacterial growth was measured three days later by selecting *Pto*DC3000 on CFC selection plats. Error bars represent standard deviations of n=6 biological replicates. ANOVA test (***, P<0.001). **(C)** Transient expression of GSTs does not impact *P. infestans* growth. Agroinfiltrated leaves expressing empty vector (EV), FLS3, GSTF2-GFP, GSTF6-GST or GSTU10-GFP were droplet-inoculated with zoospores of *P. infestans* at 50,000 spores/m. The lesion sizes were measured seven days post-infection. Error bars represent standard deviation of n=24 replicates. ANOVA test (ns, non-significant).

## DISCUSSION

In this manuscript we established photoaffinity labeling of GSTs in various plant species and use this to show safener-induced GST activities in wheat and biotic stress- and BTH-induced GSTs in Arabidopsis, independently from NPR1.

### Photoaffinity labeling of GSTs

Photoaffinity labeling is a robust way to study GSTs in plants. Conceptually, every glutathione binding protein will be labeled with DB478. In reality we only detected labeling of several GSTs from the Tau(U) and Phi(F) classes, consistent with the 23-24 kDa signals displayed on fluorescent protein gels. Labeling only Tau(U) and Phi(F) classes is not surprising because these are the two most abundant GST classes in plants (Cummins et al., 2011). Labeling of GFP/His-tagged GSTs upon overexpression by agroinfiltration confirms that these proteins are labeled with DB478.

A strong correlation between GST protein abundance and DB478 labeling indicates that DB478 is not selective to specific GSTs, but will label diverse glutathione conjugating GSTs in extracts. These observations are consistent with the detection of family-wide labeling of mammalian GSTs representing alpha, mu, pi, kappa, theta and zeta classes (Stoddard et al., 2017). Probe-enriched proteins are almost exclusively GSTs, demonstrating that this photoaffinity probe is highly selective. Although we detected several other proteins in pull-down assays using mass spectrometry, these are also detected in the no-probe control and seem to be caused by nonselective click chemistry reactions, consistent with the higher background of the ‘click only’ control in fluorescent gels.

A similar set of GSTs has been identified with pulldown assays using glutathione beads (Sappl et al., 2004; Dixon & Edwards, 2010), but these assays involve a longer purification protocol and require much more starting material. By contrast, photoaffinity labeling of GSTs can be performed on 10 μg protein scale, without purification, following a simple, 3-hour protocol, which allowed us to test and compare more conditions, treatments and mutants. We also demonstrated that GST photoaffinity labeling can be applied to a broad range of plant species, using examples of Arabidopsis, tomato, maize, wheat, liverwort and moss. This procedure can be applied to address a wide range of research questions, illustrated by our study of GST activation upon safener treatment in wheat and upon biotic stress in Arabidopsis.

### Safener activates GSTs in wheat

Using photoaffinity labeling of GSTs, we demonstrated that GSTs are activated in wheat seedlings upon treatment with safener mefenpyr-diethyl. Likewise, safener fenclorim transcriptionally induced five GSTs (U25, U24, U19, U4, and U3) in Arabidopsis (Skipsey et al., 2011), and various safeners caused GSTU19 protein accumulation in Arabidopsis (Deridder et al., 2002). Interestingly, cotreatment of safeners with herbicide seems to induce GST levels beyond that of safener alone. Although this possible synergism is not significant in our assays, this phenomenon is not much studied in plants. By contrast, the synergistic and antagonistic effects of simultaneous drug application on drug metabolizing enzymes have been extensively studied in mammals before (Borisy et al., 2003; Roell et al., 2017; Tallarida, 2001). It is interesting to note that the molecular mechanism(s) underpinning the transcriptional activation of genes encoding metabolic enzymes by safeners and herbicides have not yet been resolved.

### Role of GSTs in immunity

Using photoaffinity labeling and purification of the labeled proteins, we identified six BTH-induced GSTs from Arabidopsis leaves, which are largely consistent with SA-induced GST accumulation in Arabidopsis cell cultures (Sappl et al., 2004). GSTs might act in plant defense by detoxifying pathogen-derived phytotoxins and lipid hydroperoxides produced by peroxidation of membranes (Gullner et al., 2018). Redundancy within the GST family is often thought to hamper our understanding of the role of GSTs in immunity. Exceptions are that silencing of *NbGSTU1* in *N. benthamiana* increased susceptibility to the fungal pathogen *Colletotrichum destructivum* (Dean et al., 2005), and the *gstf9* mutant of Arabidopsis was more susceptible for the fungus *Verticilium dahliae* (Gong et al., 2018). By contrast, GST overexpression can overcome the redundancy problem with examples being the overexpression of cotton *Ga*GSTF9 in Arabidopsis increasing resistance to Verticillium wilt (Gong et al., 2018) and overexpression of *Lr*GSTU5 of royal lilly in tobacco increasing resistance to *Fusarium oxysporum* (Han et al., 2016). Unfortunately, however, overexpression of our BTH-induced GSTs did not cause phenotypes in infection assays with *Pseudomonas syringae* and *Phytophthora infestans* (**Figure 6**). The absence of phenotypes might be because these GSTs might not have relevant substrates in these interactions or GST function is disturbed by the C-terminal GFP protein.

### BTH-induced GST labeling is independent of NPR1

Our observation that avirulent bacteria and BTH induces GST labeling in Arabidopsis is consistent with the observation that SA induces accumulation of GSTs proteins in Arabidopsis cell cultures, which correlates with their transcriptional activation (Sappl et al., 2004). That BTH-induced GST labeling also occurs in the *npr1* mutant is counterintuitive because BTH activates pathogenesis-related (PR) genes via NPR1. However, our observation that NPR1 is dispensable for BTH-induced GST activation is consistent with the literature. NPR1 directly binds TGA transcription factors that bind *as-1/ocs* elements in promoters of late SA-responsive genes (Kumar et al., 2022). Also many GST-encoding genes, including *GSTF8*, *GSTU7* and *GSTU19*, have *as-1/ocs* elements in their promoter that are essential for induction by SA (Zhang & Singh, 1994; Chen & Singh, 1999; Blanco et al., 2005). But GSTs are early SA-responsive genes and these are also activated without NPR1. For instance, SA-induced expression of *GSTF9 (GST6)* and *GSTU7 (GST25)* does still occur in the *npr1* mutant, even though SA-induced expression of *PR1* is absent (Uquillas et al., 2004; Blanco et al., 2005). Likewise, transcriptional activation of the promoter of maize *Zm*GSTL1 by safeners in transgenic Arabidopsis was SA dependent and was blocked in the quadrupole *tga2/3/5/6* mutant, but unaltered in the *npr1* mutant (Behringer et al., 2011). Collectively, these data indicate that whilst activation of late SA-responsive genes is dependent on NPR1, early SA-responsive genes such as GSTs are activated via TGA transcription factors independently of NPR1. Studies on the activation of GST-encoding *GNT35* in tobacco has indicated a role for oxidative signaling because *GNT35* induction by SA can be blocked with antioxidants, and oxidative stress can induce GST expression in the absence of SA (Garreton et al., 2002).

Further elucidation of the GST activation pathway will increase our understanding of how safeners induce herbicide resistance and how pathogens induce the detoxification machinery. Photoaffinity labeling of GSTs can be a helpful new instrument in further investigations since it reports on global GST activation and can be used on any plant species upon treatments with agrochemicals and pathogens.

## CONCLUSION

We have established a simple and robust assay to detect GSTs with photoaffinity labeling in extracts of various plant species and shown that this method can be used to detect GST activation upon agrochemical treatment and pathogen infection. We have used proteomics to show the labeling of several Tau- and Phi-class GSTs and identified BTH-induced GSTs, which we confirmed by labeling transiently expressed tagged GSTs. We confirm that GST activation by SA analog BTH is independent of NPR1 and show that overexpression of defence-related GSTs does not affect immunity in transient disease assays.

## METHODS

### Chemical probes, inhibitors and agrochemicals

GST inhibitors and agrochemicals were purchased from Sigma-Aldrich: ethacrynic acid (Sigma-Aldrich, SML1083), S-hexylglutathione (Sigma-Aldrich, H6886), mefenpyr-diethyl (Supelco, 46302) and fenoxaprop-P-ethyl (Supelco, 36851). DB478 was resynthesized as described before (Stoddard et al., 2017) with minor modifications (see **Supplemental Methods**). Briefly, (tert-butoxycarbonyl)-L-phenylalanine was subjected to pentaflurophenyl trifluoroacetate to generate the respective pentafluorophenyl ester *in situ*, which was reacted with propargyl amine to give amide DB475 in 83% yield. The Boc-protecting group was removed and the free amine of the resulting DB476 was acylated with bromoacetyl bromide in the presence of excess triethylamine to produce the α-bromoacetamide DB477 in 54% yield over both steps combined. The reaction of DB477 with reduced glutathione gave DB478 in 99% yield. Aliquots of DB478 are available upon request.

### Plant materials, growth conditions and agrochemical treatments

Arabidopsis (*Arabidopsis thaliana*) ecotype Columbia plants were grown on soil at 25°C at short day conditions (8/16 h light/dark regime) in a growth chamber. *Nicotiana benthamiana*, maize (*Zea mays*), tobacco (*Nicotiana tabacum*) and tomato (*Solanum lycopersicum*) were grown in soil at standard greenhouse conditions. Liverwort (*Marchantia polymorpha*) and moss (*Physcomitrella patens*) were grown under sterile conditions as described by (Thamm et al., 2020) and (Moody et al., 2021), respectively. For the BTH treatment, 5-week-old Arabidopsis plants were grown on soil in a controlled environment (short day conditions, 25°C). BION granules (Syngenta) were dissolved in water to a final concentration of 1 μM BTH and Arabidopsis plants were watered with or without BTH over three consecutive days. Arabidopsis leaf tissue was harvested on day four and frozen in liquid nitrogen till further use. Winter wheat (*Triticum aestivum*) seeds were sterilized with 30% ethanol for 5 min, 30% bleach for 10 min and rinsed six times with sterile water. Sterilized seeds were plated in Murashige and Skoog agar plates (half strength Murashige and Skoog basal [Duchefa-Biochemie, M0222] salt mixture containing 0.8% agar) and stratified at 4°C for 1 day. For germination, seeds were placed at 25°C under long day conditions (16/8 h light/dark regime). After 4 days, germinated wheat seedlings were transferred into Hoagland’s liquid medium (half strength Hoagland’s No.2 Basal Salt Mixture [Sigma-Aldrich, H2395]). After one day adaptation, seedlings were treated with 100 μM mefenpyr-diethyl; fenoxaprop-P-ethyl; mefenpyr-diethyl; or fenoxaprop-P-ethyl or an equal amount of DMSO solvent. Seedling tissue was harvested 24h after treatment and frozen in liquid nitrogen until used.

### Cloning

Used and generated plasmids are summarized in Supplemental **Table S1**. The primers used for sequence amplification are summarized in Supplemental **Table S2**. Golden Gate Modular Cloning kit (Weber et al., 2011) and Golden Gate Modular Cloning Toolbox for plants (Engler et al., 2014) were used for cloning. The binary vector pJK268c containing the P19 silencing inhibitor (pL1V2-P19-F2) was used as binary vector (Kourelis et al., 2020). The sequences encoding Arabidopsis GSTs (GSTF2, GSTU4, GSTU10, GSTF6 and GSTF7) were amplified from Arabidopsis complementary DNA (cDNA) using the forward and reverse primer pairs summarized in Supplemental **Table S2**. Using BsaI restriction sites, PCR products were combined with GG1-55, GG1-57, GG1-70 or GG1-78 (Supplemental **Table S1**) to assemble the corresponding expression constructs. Using BpiI restriction sites, PCR products were first cloned into the level-0 module GG1-01 and later transferred to pJK268c together with GG1-55, GG1-57, GG1-70 or GG1-78 in a BsaI ligation reaction. Generated expression constructs were transformed into *E. coli* for their amplification and sequencing. Afterwards, constructs were transformed into Agrobacterium (*Agrobacterium tumefaciens*) strain GV3101, carrying the pMP90 helper plasmid. Agrobacterium transformants were selected on Lysogeny Broth (LB) agar plates (10 g/L NaCl, 10 g/L tryptone, 5 g/L yeast extract, 15 g/L agar) containing 50 μM gentamycin, 25 μM rifampicin and 50 μM kanamycin. Single colonies were picked and grown in liquid LB (10 g/L NaCl, 10 g/L tryptone, 5 g/L yeast extract) containing the same antibiotic used before. Glycerol stocks of the selected transformants were prepared mixing Agrobacterium cultures with 50% sterile glycerol in a 1:1 ratio.

### Agroinfiltrations

*N. benthamiana* plants were grown on soil at greenhouse conditions described before. *A. tumefaciens* GV3101-pMP90 carrying the desired expression constructs were grown overnight (approximately 18 h) at 28°C with agitation in LB media containing 50 μM gentamycin, 25 μM rifampicin and 50 μM kanamycin. Bacterial cells were collected by centrifugation at 1000 x *g* for 10 min at room temperature, washed with infiltration buffer (10 mM MES, 10 mM MgCl_2_, pH 5.6 and 150 μM acetosyringone), diluted to an optical density at 600 nm (OD600) of 0.5. Bacteria were incubated at 28°C for 2 h and afterwards, the two new fully-expanded leaves of 4-5 week old *N. benthamiana* plants were hand-infiltrated with the bacterial suspensions with the help of a needle-less 10 ml syringe. Agroinfiltrated *N. benthamiana* leaf tissue was collected at 4 days post-infiltration (dpi) or used for infections assays. For *P. infestans* infection assays, *N. benthamiana* leaves were half infiltered with the empty vector (EV) control and half infiltrated with the construct of interest (GST expressing construct).

### Protein extraction

Leaf tissue from various plant species was frozen in liquid nitrogen and ground to fine powder with a mortar and pestle. Arabidopsis total proteins were extracted in cold homogenization buffer (100 mM Tris-HCl pH 7.4) in a 1:3 weight-to-volume ratio. For the other plant species, total proteins were extracted in cold Tris-HCl buffer containing 3 mM β-mercaptoethanol (βME). The extracts were incubated for 20 min rotating at 4°C for complete homogenization and centrifuged at 5,000 x *g* for 20 min at 4°C. Supernatants containing soluble proteins were filtered through a layer of miracloth filter and centrifuged again at 5,000 x *g* for 20 min at 4°C. Frozen wheat seedlings were ground to fine powder with the help of a tissue lyser (QIAGEN, TissueLyser II) in 1.5 mL safe-lock Eppendorf tubs containing three 2.4 mm metal beads. Next, total proteins were extracted in cold Tris-HCl homogenization buffer in a 1:3 weight-to-volume ratio and cleared by centrifugation at 10,000 x *g* for 10 min at 4°C. For all extractions, total protein concentration was determined by DC protein assay Kit (Bio-Rad, 5000116) with bovine serum albumin (BSA) standards. Protein concentration was adjusted to 0.5 mg/mL for each sample and the resulting samples were used for labelling without freeze thawing.

### Labeling and click chemistry

DB478 was prepared at 50 - 1000 μM stock solutions in dimethyl sulfoxide (DMSO). Samples of 50 μl were incubated with or without DB478 at the indicated probe concentration (usually 5 μM) and left 20 min incubating before UV radiation (365 nm, in microtiter plates on ice) for the indicated amount of time (usually 45 min). Equal volumes of DMSO were added to no-probe controls. For inhibition experiments, 50 μL extracts were pre-incubated with competitor molecules, 0.2 mM of S-hexylglutathione (SHG) or ethacrynic acid (EA), for 15 min before DB478 labelling. Labelling reactions were stopped by adding 1:4 (v/v) cold acetone and precipitated by 3 min centrifugation at maximum speed using a benchtop centrifuge. Protein pellets were resuspended in 44.5 μl of PBS-SDS buffer (phosphate-buffered saline (PBS) pH 7.4 with 1% (w/v) sodium dodecyl sulfate (SDS)) and heated for 3 min at 90°C. For click chemistry, 44.5 μL of labelled proteins were incubated with 2 μM Cy5 Picolyl Azide (Click Chemistry Tools, 50 μM stock in DMSO) or Fluorescein Picolyl Azide (Click Chemistry Tools, 50 μM stock in DMSO) and a premixture of 100 μM tris((1-benzyl-1H-1,2,3-triazol-4-yl) methyl)amine (TBTA; 3.4 mM stock in DMSO:t-butanol 1:4), 2 mM tris(2-carboxyethyl)phosphine hydrochloride (TCEP; 100 mM stock in water) and 1 mM copper(II) sulfate (CuSO_4_; 50 mM stock in water). Samples were incubated for 1 h at room temperature in the dark and quenched by acetone precipitation. Protein pellets were resuspended in 50 μL of 2x gel loading buffer (100 mM Tris-HCl pH 6.8, 200 mM dithiotreitol, 4% SDS, 20% glycerol and 0.02% bromophenol blue) and heated for 5 min at 90°C.

### In-gel fluorescence scanning and western blot

Labelled proteins were loaded and separated on 10-12% SDS-PAGE acrylamide gels at 150 V. Fluorescence scanning of acrylamide gels was performed on an Amersham Typhoon 5 scanner (GE Healthcare Life Sciences) using excitation and emission wavelengths of ex488/em520 and ex635/em670 for Cy2 and Cy5 scanning, respectively. Scanned gels were stained with Coomassie Brilliant Blue G-250. For Western Blot, proteins separated in SDS-PAGE gels were transferred to a PVDF membrane using Trans-Blot Turbo Transfer System (Bio-Rad). For anti-His and anti-GFP blots, membranes were blocked in 5% (w/v) milk in TBS-T (50 mM Tris-HCl pH 7.6, 150 mM NaCl and 0.1% Tween-20) overnight at 4°C and incubated in 1:5000 anti-GFP-HRP (Abcam, ab6663) or anti-6xHis-HRP (Invitrogen, #R931-25) in TBS-T/milk 5% for 1.5 h at room temperature. For anti-PR2 blots, membranes were blocked in TBS-T/milk 5% overnight at 4°C and incubated in 1:2000 anti-PR2 (Agrisera, AS122366) in TBS-T/milk 5% for 1.5h at room temperature. Membranes were washed three times for 5 min in TBS-T at room temperature and incubated in 1:10000 secondary anti-Rabbit-HRP (Invitrogen, #31460) in TBS-T/Milk 5% for 1 h at room temperature. Before scanning, all blots were washed six times for 5 min in TBS-T/PBS-T and a last wash for 5 min in TBS/PBS. Clarity Western ECL substrate (Bio-Rad) was used for chemiluminescent protein detection.

### Large-scale labelling and affinity purification

A volume of 1 mL protein extracts (0.5 mg/mL protein concentration) were incubated with or without 5 μM DB478 as described in “Labeling and click chemistry”. Labelling reactions were stopped by methanol/chloroform precipitation (addition of 4 volumes of ice-cold methanol, 1 volume of ice-cold chloroform and 3 volumes of water) and precipitated by 40 min centrifugation at 4,000 x *g* at 4°C. Protein pellets were resuspended in 1 mL of PBS-SDS buffer by bath sonication and heated at 90°C for 5 min. Labelled proteins were biotinylated with 20 μM azide-PEG3-biotin (Sigma-Aldrich, 2 mM stock in DMSO) using click chemistry as described in “Labeling and click chemistry”. Samples were incubated for 1 h at room temperature in the dark, quenched by the addition of 10 mM ethylenediaminetetraacetic acid (EDTA) and precipitated via the methanol/chloroform method. Protein pellets were resuspended in 1.5 ml of 1.2% (w/v) SDS dissolved in PBS by bath sonication and diluted by adding 7.5 ml of PBS. The resulting solution was incubated with 100 μL of pre-equilibrated Pierce High Capacity Streptavidin Agarose beads (Thermo Fisher Scientific, 10302384) for 2h at room temperature. Agarose beads containing the labelled proteins were collected by centrifugation at 1,000 x *g* for 5 min at room temperature. Agarose beads were washed three times with 10 mL of PBS-1% SDS buffer, three times with PBS and a final wash with 10 mL of water.

### On-bead trypsin and Lys-C digestion

For on-bead digestion, the captured agarose beads containing the labelled proteins were treated with 500 μL 6 M urea dissolved in PBS and 10 mM TCEP (200 mM stock in water) for 15 min at 65°C. A final concentration of 20 mM 2-chloroacetamide (CA, 400 mM stock in water) was added and the sample incubated for 30 min at 35°C in the dark. The reaction was diluted by the addition of 950 μL of PBS and the supernatant was removed by centrifuging the beads at 500 g for 2 min. 100 μL of Lys-C digestion solution (Walko, 5 μg of Lys-C were reconstituted in 100 μL 1 M urea/ 50 mM Tris-HCl buffer pH 8) was added to the beads and left incubating at 37°C for 3 h. 100 μL of trypsin digestion solution (Promega, 5 μl of reconstituted trypsin were dissolved in 100 μL 50 mM Tris-HCl buffer pH 8) was added to the beads and left incubating at 37°C overnight. The supernatant containing the digested peptides was dried by vacuum centrifugation and submitted for MS analysis.

### LC-MS/MS and peptide identification using MaxQuant

MS samples were analyzed as described in (Morimoto et. al., 2019). Briefly, the acidified tryptic peptides were desalted using C18 Tips (Thermo Fisher Scientific, 10 μL bed, 87782) and peptide samples were resuspended in 0.1% FA solution and analysed on an Orbitrap Elite instrument (ThermoFisher Scientific) (Michalski et al., 2012) attached to an EASY-nLC 1000 liquid chromatography system (Thermo Fisher Scientific). Samples were separated on an analytical column based on a fused silica capillary with an integrated PicoFrit emitter (New Objective) with Reprosil-Pur 120 C18-AQ 1.9 μm resin. Peptides were next separated using 140 min gradient of solvent and analysed with Orbitrap analyser (using Fourier Transform MS) at a resolution of 60,000 with the internal lock-mass option switched on (Olsen et al., 2005).

RAW spectra were submitted to an Andromeda search in MaxQuant (version 1.5.3.30) using the default settings (Cox et al., 2011; Cox and Mann, 2008). Label-free quantification and match between-runs were activated (Cox et al., 2014). The MS/MS spectra data were searched against the UniProt Mus musculus reference database (55,412 entries) and the UniProt *Arabidopsis thaliana* reference database (39,359 entries). All searches included a contaminants database (as implemented in MaxQuant, 245 entries). Enzyme specificity was set to ‘Trypsin/P’ and the MS/MS match tolerance was set to ±0.5 Da. The peptide spectrum match FDR and the protein FDR were set to 0.01 (based on target-decoy approach) and the minimum peptide length was seven amino acids. For protein quantification, unique and razor peptides were allowed. Modified peptides were allowed for quantification with a minimum score of 40. Relative quantification of proteins between different MS runs is based exclusively on the label-free quantifications as calculated by the software MaxQuant, using the “MaxLFQ” algorithm (Cox et al., 2014). Filtering of results was done in Perseus version 1.6.13.0 (Tyanova et al., 2016). Briefly, data was filtered against contaminants and only those proteins found in at least one group in all three replicates were considered. Further analysis and graphs were performed using the Qfeatures and ggplot R packages (Gatto, 2021). Imputed values were generated using a missing not at random (MNAR) and missing at random (MAR) mixed imputation over the whole matrix, and the fold change and adjusted P values (adj. P Val) were calculated using the fdr method over the three biological replicates. The correct clustering of the biological replicates by categorical groups was evaluated using principal component analysis (PCA) plots.

### Disease assays

Agromonas assays with *Pto*DC3000 infections were performed as described in (Buscaill et al., 2021) with some modifications. Briefly, *Pto*DC3000 was grown from glycerol in LB containing 25 μg/mL rifampicin overnight. Overnight culture was centrifugated at 1500 x *g* for 10 min at room temperature, washed one time with sterile water and resuspended to a density of OD600=1. Bacterial suspension was sprayed on the adaxial leaf surface of agroinfiltrated leaves of *N. benthamiana*. Previous to bacterial infection, plants were covered with a humidity dome for a day. Once sprayed, *N. benthamiana* plants were re-covered with the humidity dome and left for 3 days at 21°C under long day conditions (16/8 h light/dark regime) in a growth chamber. For bacterial growth calculations, three leaf discs (1 cm diameter) per plant were excised from the treated leaves and surface sterilised in 15% H_2_O_2_ for 2 min. Leaf disc were rinsed twice with sterile water and dried under sterile conditions. Leaf discs were ground in 1 mL sterile water with the help of a tissue lyser (QIAGEN, TissueLyser II) using 1.5 mL safe-lock Eppendorf tubs containing three 2.4 mm metal beads. A serial dilution of the leaf homogenate was plated onto LB agar plates containing cetrimide, fucidin and cephaloridine (CFC supplement, OXOID, OXSR0103E) for selection of *P. syringae* but not *A. tumefaciens* colonies. Plates were incubated at 28°C for 3 days and *Pto*DC3000 colonies where counted using the best dilution plated. *Pto*DC3000 bacterial growth is expressed in the log_10_ transformation of the number of colony-forming unit (CFU) per cm^2^ of leaf tissue (log_10_CFU/cm^2^). *P. infestans* (strain 88069) infections were performed as described in (Song et al., 2009) with some modifications. Briefly, *P. infestans* was grown on rye sucrose agar medium at 18°C under dark conditions for 2 weeks. Zoospores were collected by the addition of cold sterile water into the plates and its incubation at 4°C for a minimum of 4 h. Spore concentration was measured with the help of a haemocytometer and was adjusted to 1×10^5^ zoospores/mL. Three days after agroinfiltration, *N. benthamiana* leaves were detached and infected by drop-inoculation of 10 μL of the zoospores culture on the abaxial leaf surface. Infected leaves were incubated in moist trays at 18°C for 7 days. The size of the infection area was measured by scanning the leaves at ex635/em670 and measuring the lesion area by ImageJ quantification. The successful expression of the GFP-tagged GST expression construct was detected by scanning the leaves at ex488/em520. Arabidopsis plants were infected by infiltrating bacterial suspensions at OD600 = 0.005 for the infiltration with syringe without needle. Three to six alternating leaves of a single plant were infiltrated sufficiently with one strain. Ultra-pure water was used for the infiltration of the control plants. Half of the plants were used for the protein extraction two days post-infection (2dpi) and the other half for a phenotypical analyses (5dpi).

### Image quantification and statistical analysis

ImageJ (U.S. National Institutes of Health) was used for the quantification of the fluorescence of the labelled proteins and *P. infestans* lesions. To quantify fluorescence signals from protein gels, scanned images (.gel format) were used. The intensity of the defined signal area was quantified using the Analyse -> Measure tool. The integrated density (IntDen) was used to compare intensities signal of the same dimensions (area) measurements within the same protein gel. To quantify *P. infestans* infection lesions, leaf scans were calibrated using the Analyse -> Set scale tool, using a ruler scan for calibration. All images to be compared were set to the same format (8-bit jpg) and an automatic threshold was applied (Image -> Adjust -> Threshold). Set areas applied by the threshold were measured using the Analyse -> Measure tool. The area measured in cm^2^ was used to compare lesion sizes between GST expressing constructs and the EV control. All disease score data and fluorescence quantifications were subjected to statistical analysis using RStudio software. P-values were calculated using one-way analysis of variance (on-way ANOVA). All values shown are mean values, and the error intervals represent the standard deviation (sd).

## DECLARATIONS

### Ethics approval and consent to participate

not applicable

### Consent for publication

not applicable

### Availability of data and materials

All data generated or analysed during this study are included in this published article and its supplementary information file.

### Competing interests

The authors declare that they have no competing interests

### Funding

This project was financially supported by BBSRC iCASE studentships with Syngenta (project DDT00060, MFF, DB); ERG-CoG-2013 project 616449 ‘GreenProteases’ (RH); and ERC-AdG-2019 project 101019324 ‘ExtraImmune’ (RH).

### Authors’ contributions

MFF designed and performed most of the experiments; ATW and BJK provided the initial probes; DB resynthesized the probes under guidance of JB; FK and MK performed LC-MS/MS analysis; RvdH conceived and supervised the project; MFF and RvdH wrote the manuscript together with input from all authors. The funding body had no influence on the design of the study and collection, analysis, and interpretation of data and in writing the manuscript.

## Acknowledgements

We like to thank Urszula Pyzio for excellent plant care and Sarah Rodgers and Caroline O’Brien for excellent technical support. We thank Steven Spoel for providing the *npr1-1* mutant, Svenja Heimann and Jenny Bormann for proteomics support, and Alexandra Casey and Laura Moody for growing liverwort and Physcomitrella, respectively.

## Notes

### Competing Interest Statement

The authors have declared no competing interest.

### Summary of Updates

Corrected surname of co-author.

