## Supplemental Table for "Glutathione transferase photoaffinity labeling demonstrates GST activation by safeners and NPR1-independent activation by BTH"

### Supplemental Tables

**Table S1** Used plasmids

| Plasmid | Description | Reference |
|---|---|---|
| pJK268c | pL1V2-P19-F2, Binary vector. | Kourelis et al., 2020 |
| pICH41414 | pL0M-T-35S-1-41414, Level 0 Module, 35S terminator, Golden Gate plant kit | Engler et al., 2014 |
| pICH51288 | pL0M-PU-35S-TMV-3-51288, Level 0 Module, 2 x 35S promoter + TMV Ω (0.8 kb), Golden Gate plant kit | Engler et al., 2014 |
| EC15456 | pL0V-SC1-15456, Level 0 Cloning vector, Golden Gate original vector | |
| EC15259 | pL0M-C2-6xHis-15259, Level 0 Module, His for C-terminal fusion | |
| EC15095 | pL0M-C2-eGFP-15095, Level 0 Module, eGFP for C-terminal fusion | |
| pMF358 | pL2M-P19-kan-2x35S::AtGSTF2-HIS | This work |
| pMF359 | pL2M-P19-kan-2x35S::AtGSTF2-GFP | This work |
| pMF360 | pL2M-P19-kan-2x35S::AtGSTU4-HIS | This work |
| pMF361 | pL2M-P19-kan-2x35S::AtGSTU4-GFP | This work |
| pMF364 | pL2M-P19-kan-2x35S::AtGSTU10-HIS | This work |
| pMF365 | pL2M-P19-kan-2x35S::AtGSTU10-GFP | This work |
| pMF366 | pL2M-P19-kan-2x35S::AtGSTF6-HIS | This work |
| pMF367 | pL2M-P19-kan-2x35S::AtGSTF6-GFP | This work |
| pMF368 | pL2M-P19-kan-2x35S::AtGSTF7-HIS | This work |
| pMF369 | pL2M-P19-kan-2x35S::AtGSTF7-GFP | This work |
| pJK668 | pL2M-kan-2x35S::FLS3 | Buscaill et al., 2021 |

**Table S2** Used oligonucleotides

| Name | Sequence (5′ to 3′)* |
|---|---|
| GSTU10f | **TTGAAGACAA**AATGGAGGAGAAGAAGAGCAAAG |
| GSTU10r | **TTGAAGACAA**CACCTGCATTTGCAGCCTGC |
| GSTF2f | TTGGTCTCAAATGGCAGGTATCAAAGT |
| GSTF2r | TTGGTCTCAGTCACCATCTTCAAAGGC |
| GSTF2f | TTGGTCTCATGACCTCAAGCTCTTCGAATC |
| RV2_GSTF2r | TTGGTCTCACACCCTGAACCTTCTCGGAAG |
| FW_GSTF6f | TTGGTCTCAAATGGCAGGAATCAAAG |
| RV_GSTF6r | TTGGTCTCACACCAAGAACCTTCTGAGCAG |
| FW_GSTF7f | TTGGTCTCAAATGGCAGGAATCAAAG |
| RV_GSTF7r | TTGGTCTCACACCAAGAACCTTCTTAGCAG |
| FW_GSTU8 | TTGGTCTCAAATGAACCAAGAAGAGCACG |
| RV_GSTU8r | TTGGTCTCACACCATTAGATGTAACACTTC |

*, Highlighted are the BsaI restriction sites (underlined) and the BpiI restriction sites (bold).
