## Supplemental Methods for "Glutathione transferase photoaffinity labeling demonstrates GST activation by safeners and NPR1-independent activation by BTH"

**Supplemental Methods:** Synthesis of DB478

### General experimental procedures

***NMR spectra:*** Proton ($^1$H) and carbon ($^{13}$C) NMR spectra were recorded on a Bruker AV 600 (600/126 MHz), Bruker AV 500 (500/125 MHz), Bruker AV 400 (400/100 MHz) or Bruker DPX 200 (200/50 MHz) spectrometer. Proton and carbon chemical shifts are quoted in ppm and referenced to residual protonated solvent. Assignments were made on the basis of chemical shifts, coupling constants, COSY, HSQC, HMBC data and comparison with spectra of related compounds. Resonances are described as s (singlet), d (doublet), t (triplet), q (quartet), sept (septet), m (multiplet), br (broad), dd (double doublet) and so on. Coupling constants (*J*) are given in Hz and are rounded to the nearest 0.5 Hz. Diastereotopic protons and carbons are distinguished with (‘) for example CHH’ refers to a $CH_2$ group with diastereotopic protons.

***Mass spectra:*** Low resolution mass spectra were recorded on a Fisons Platform spectrometer (ES). High resolution mass spectra were recorded by mass spectrometry staff at the Chemistry Research Laboratory, University of Oxford, using a Bruker Daltronics microTOF spectrometer (ES) or a Micromass GCT (FI). m/z values are reported in Daltons with their percentage abundances and, where known, the relevant fragment ions in parentheses. High resolution values are calculated to four decimal places from the molecular formula, all found values being within a tolerance of 5 ppm.

***Infrared spectra*****:** Infrared spectra were recorded on a Bruker Tensor 27 Fourier Transform spectrometer, using diamond ATR. Absorption maxima ($\nu_{max}$) are quoted in wavenumbers (cm$^{-1}$).

***Optical rotation*****:** Optical rotations were measured using a Perkin-Elmer 241 polarimeter in a cell of 1 dm path length (l). Specific rotations denoted as $[\alpha]_D^T$ where T = temperature in °C, D refers to the 145 line of a sodium lamp (λ = 589 nm), and are calculated according to the equation below where α is the observed rotation: $[\alpha]_D^T = \frac{100a}{lc}$.

***Chromatography techniques:*** TLC was performed on Merck DC-Alufolien 60 F254 0.2 mm precoated plates and visualised using an acidic vanillin or basic potassium permanganate dips. Flash column chromatography was performed on Merck 60 silica (particle size 40–63 μm, pore diameter 60 Å) and the solvent system used is recorded in parentheses.

***Reactions:*** All non-aqueous reactions were carried out in flame dried glassware under an inert atmosphere of nitrogen and employing standard techniques for handling air-sensitive materials unless specified otherwise. Solvents and commercially available reagents were dried and purified before use, as appropriate. All water used experimentally was distilled and the term ‘brine’ refers to a saturated solution of sodium chloride in water.

### Synthesis of DB475

*tert*-Butyl *(S)*-(3-(4-benzoylphenyl)-1-oxo-1-(prop-2-yn-1-ylamino)propan-2-yl)carbamate

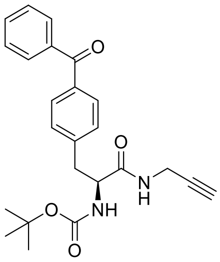

Pentafluorophenyl trifluoroacetate (0.22 mL, 1.3 mmol, 1.3 equiv.) was added dropwise to a stirred solution of (*tert*-butoxycarbonyl)-L-phenylalanine (0.37 g, 1.0 mmol, 1.0 equiv.) and pyridine (0.097 mL, 1.2 mmol, 1.2 equiv.) in DMF (10 mL) and the reaction mixture was stirred at room temperature for 1 h. The reaction mixture was diluted with EtOAc (20 mL) and then washed with sat. $NaHCO_3$ solution (20 mL), and brine (3 × 30 mL), dried ($MgSO_4$), filtered and solvent removed *in vacuo* to give the crude pentafluorophenyl ester which was passed through a short silica plug (EtOAc) to remove by- products. The crude product was redissolved in DMF (5 mL), Propargyl amine (0.058 mL, 1.0 mmol, 1.0 equiv.) and DIPEA (0.16 mL, 1.1 mmol, 1.1 equiv.) were added and then the reaction mixture was stirred at rt for 3 h. The reaction mixture was diluted with EtOAc (20 mL), washed with 1 M HCl (20 mL), sat. $NaHCO_3$ (20 mL), and brine (3 × 20 mL), dried ($MgSO_4$), filtered, and solvent removed *in vacuo* to give the title compound as a yellow oil (0.25 g, 0.83 mmol, 83%) which required no further purification. $^1$H NMR (400 MHz, $CDCl_3$) δ 7.83 – 7.71 (m, 4H, 4 × ArC**H**), 7.64 – 7.55 (m, 1H, ArC**H**), 7.52 – 7.44 (m, 2H, 2 × ArC**H**), 7.38 – 7.29 (m, 2H, ArC**H**), 6.15 (s, 1H, N**H**), 5.01 (s, 1H, C**H**), 4.39 (d, *J* = 8.0 Hz, 1H, C**H**$NHCO_2C(CH_3)_3$), 4.05 – 3.94 (m, 2H, NHC**H₂**C≡CH), 3.17 (qd, *J* = 14.0, 7.0 Hz, 2H, C**H₂**$CHNHCO_2C(CH_3)_3$), 2.21 (t, *J* = 2.5 Hz, 1H, $NHCH_2C$≡C**H**), 1.42 (s, 9H, $CHNHCO_2C$(C**H₃**)$_3$); LRMS *m/z* (ESI$^+$) 429.2 (M+Na$^+$, 100%). These data are consistent with the literature (Stoddard et al., 2017).

### Synthesis of DB476

*(S)*-2-Amino-3-(4-benzoylphenyl)-*N*-(prop-2-yn-1-yl)propenamide

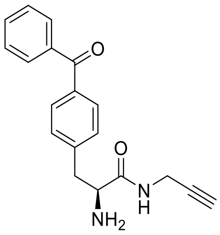

Trifluoroacetic acid (0.64 mL, 8.3 mmol, 10 equiv.) was added to a stirred solution of **DB475** (0.25 g, 0.83 mmol, 1.0 equiv.) and the reaction mixture was stirred at rt for 18 h. Solvent was removed *in vacuo* to give the title compound as a yellow oil (0.20 g, 0.83 mmol, 100%) which required no further purification. $^1$H NMR (400 MHz, DMSO-$d_6$) δ 8.84 (t, *J* = 5.5 Hz, 1H, ArC**H**), 8.16 (d, *J* = 5.0 Hz, 2H, N**H**), 7.64 (ddt, *J* = 9.5, 6.5, 1.5 Hz, 4H, 4 × ArC**H**), 7.54 – 7.45 (m, 2H, 2 × ArC**H**), 7.39 – 7.31 (m, 2H, 2 × ArC**H**), 3.93 (d, *J* = 7.0 Hz, 1H, C**H**$NH_2$), 3.86 (dd, *J* = 5.5, 2.5 Hz, 2H, C**H₂**$CHNH_2$), 3.16 (t,

$J$ = 2.5 Hz, 1H, NHCH$_2$C≡C**H**), 3.12 – 2.94 (m, 2H, NHC**H$_2$**C≡CH); LRMS *m/z* (ESI$^+$) 307.0 (M+H$^+$, 100%). These data are consistent with the literature (Stoddard et al., 2017).

#### Synthesis of DB477

*(S)*-3-(4-benzoylphenyl)-2-(2-bromoacetamido)-*N*-(prop-2-yn-1-yl)propenamide

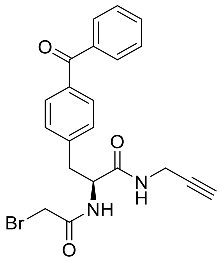

Bromoacetyl bromide (0.048 mL, 0.55 mmol, 1.1 equiv.) was added dropwise to a stirred solution of **DB476** (0.20 g, 0.50 mmol, 1.0 equiv.) and triethylamine (0.15 mL, 1.1 mmol, 2.2 equiv.) in $CH_2Cl_2$ (5 mL) at 0 °C and the reaction mixture was warmed to rt and stirred for 4 h. After this time, the solvent was removed *in vacuo* and the resulting crude residue was further purified by flash column chromatography ($CH_2Cl_2$ → 5% acetone/ $CH_2Cl_2$) to give the title compound (0.10 g, 0.26 mmol, 54%) as a white solid. $R_f$ 0.28 (5% acetone / $CH_2Cl_2$); $^1$H NMR (500 MHz, $CDCl_3$) δ 7.80 – 7.75 (m, 4H, 4 × ArC**H**), 7.63 – 7.56 (m, 1H, ArC**H**), 7.49 (dd, $J$ = 8.5, 7.0 Hz, 2H, 2 × ArC**H**), 7.40 – 7.31 (m, 2H, 2 × ArC**H**), 7.06 – 7.02 (m, 1H, N**H**), 5.90 (d, $J$ = 6.0 Hz, 1H, N**H**), 4.62 (td, $J$ = 7.5, 6.5 Hz, 1H, CH$_2$C**H**NH), 4.10 – 3.91 (m, 2H, C**H$_2$**CHNH), 3.91 – 3.82 (m, 2H, NHCO$_2$C**H$_2$**Br), 3.28 – 3.13 (m, 2H, NHC**H$_2$**C≡CH), 2.22 (t, $J$ = 2.5 Hz, 1H, NHCH$_2$C≡C**H**); $^{13}$C NMR (126 MHz, $CDCl_3$) δ 196.2 (**C**=O), 169.3 (**C**=O), 165.7 (**C**=O), 140.7 (**C**CO), 137.5 (**C**CO), 136.7 (**C**CH$_2$), 132.5 (Ar**C**H), 130.7 (Ar**C**H), 130.0 (Ar**C**H), 129.4 (Ar**C**H), 128.3 (Ar**C**H), 78.5 (C≡**C**H), 72.16 (**C**≡CH), 54.8 (NH**C**H$_2$Br), 38.2 (CH$_2$**C**HNH), 29.3 (**C**H$_2$CHNH), 28.6 (NH**C**H$_2$C≡CH); LRMS *m/z* (ESI$^+$) 449.0 (M+Na$^+$, 100%) 451.0 (M+Na$^+$, 100%); HRMS *m/z* (ESI$^+$): found 449.0470, $C_{21}H_{19}{}^{79}BrN_2O_3Na$ (M+Na$^+$) requires 449.0471; HRMS *m/z* (ESI$^+$): found 451.0450, $C_{21}H_{19}{}^{81}BrN_2O_3Na$ (M+Na$^+$) requires 451.0471.

#### Synthesis of DB478

*N$^5$*-(*(R)*-3-((2-(((*S*)-3-(4-Benzoylphenyl)-1-oxo-1-(prop-2-yn-1-ylamino)propan-2-yl)amino)-2-oxoethyl)thio)-1-((carboxymethyl)amino)-1-oxopropan-2-yl)-l-glutamine

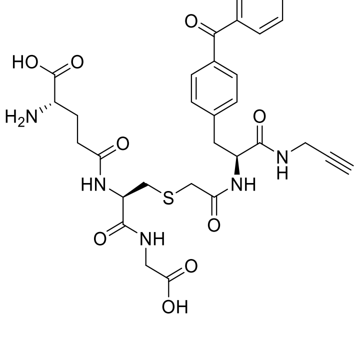

L-Glutathione reduced (0.031 g, 0.052 mmol, 1.1 equiv.) and potassium iodide (0.0040 g, 0.024 mmol, 0.50 equiv.) were added to a stirred solution of **DB477** (0.020 g, 0.047 mmol, 1 equiv.) in DMSO (0.5 mL) and the reaction mixture was stirred at room temperature for 18 h. The reaction mixture was then purified by reverse- phase preparative HPLC (out of house) to give the title compound as a white solid (0.030 g, 0.046 mmol, 99%). $^1$H NMR (400 MHz, $D_2O$) δ 7.67 (m, 5H, 5 × ArC**H**), 7.50 (m, 2H, 2 × ArC**H**), 7.36 (d, 2H, 2 × ArC**H**), 4.61 (t, *J* = 8.0 Hz, 1H, $H_2$NC**H**$CH_2$), 4.38 (t, *J* = 9.0, Hz, 1H, C**H**$CH_2$S), 3.85 (m, 5H, NHC$\mathbf{H_2}$COOH, ArC$\mathbf{H_2}$CH , NHC**H**CONH), 3.20 (m, 2H, NHC$\mathbf{H_2}$C≡CH), 3.04 (dd, *J* = 13.5, 9.0 Hz, 2H, $CH_2$C$\mathbf{H_2}$CONH), 2.74 – 2.53 (m, 2H, SC$\mathbf{H_2}$CONH), 2.49 (d, *J* = 2.5 Hz, 1H, NH$CH_2$C≡C**H**), 2.43 (t, *J* = 7.5 Hz, 2H, SC$\mathbf{H_2}$CH), 2.07 (d, J = 7.0 Hz, 2H, $CH_2$C$\mathbf{H_2}$CH$NH_2$); LRMS ($ESI^-$) 652.2 ($M-H^+$). These data are consistent with the literature (Stoddard et al., 2017).
